# Anti-adipogenic properties of clock activator chlorhexidine and a new derivative

**DOI:** 10.1101/2023.10.12.562086

**Authors:** Xuekai Xiong, Tali Kiperman, Weini Li, Zhipeng Fang, Alon Agua, Wendong Huang, David Horne, Ke Ma

## Abstract

**Background:** The circadian clock exerts temporal control of metabolic pathways to maintain homeostasis, and its disruption leads to the development of obesity and insulin resistance. In adipose tissue, key regulators of clock machinery orchestrate adipogenic processes via the Wnt signaling pathway to impact mature adipocyte development.

**Methods:** Based on the recent finding of chlorhexidine as a new clock activator, we determined its potential anti-adipogenic activities in distinct adipogenic progenitor models. Furthermore, we report the structural optimization of chlorhexidine leading to the discovery of analogs with improved efficacy in inhibiting adipogenesis.

**Results:** In adipogenic progenitors with *Per2::dLuc* luciferase reporter, Chlorhexidine shortened clock period length with induction of core clock components. Consistent with its clock-activating function, Chlorhexidine robustly suppressed the lineage commitment and maturation of adipogenic mesenchymal precursors, with comparable effect on inhibiting preadipocyte terminal differentiation. Mechanistically, we show that Chlorhexidine induces signaling components of the Wnt pathway resulting in activation of Wnt activity. Via modification of its chemical scaffold, we generated analogs of chlorhexidine that led to the identification of CM002 as a new clock-activating molecule with improved anti-adipogenic activity.

**Conclusions:** Collectively, our findings uncovered the anti-adipogenic functions of a new class of small molecule clock activators. These compounds provide novel chemical probes to dissect clock function in maintaining metabolic homeostasis and may have therapeutic implications in obesity and associated metabolic disorders.

## Introduction

Circadian clock, a molecular machinery that drives ∼24 hour oscillations, exerts pervasive temporal regulation in diverse metabolic processes (1, 2). Disruption of this timing mechanism, increasingly prevalent in our modern lifestyle, leads to metabolic dysregulations contributing to the development of obesity and Type II diabetes (3-5). Pharmacological modulation of circadian clock and its metabolic outputs may have therapeutic applications for metabolic diseases (6-8). At present, there are increasing efforts to identify clock-modulatory small molecules that may have potentials for clock-targeted therapies (9-12).

The circadian clock rhythm is generated by a transcriptional/translational feedback circuit (1) composed of positive and negative regulatory components. Transcription activators CLOCK (Circadian Locomotor Output Cycles Kaput) and Bmal1 (Brain and Arnt-Like 1) initiates clock transcription that is countered by the feedback inhibition of their direct target genes and transcription repressors, Periods (Per 1-3) and Cryptochromes (Cry1 & 2). In addition, RORα (Retinoid-related Orphan Receptor α) and Rev-erbα, a pair of antagonistic transcription factors, exerts positive and negative regulations respectively, to control Bmal1 transcription that forms an re-enforcing loop of the clock mechanism (13, 14). Interestingly, both the positive and negative arms of the core clock feedback loop modulate adipogenic process. Loss of *CLOCK* or *Bmal1* resulted in obesity (Paschos, Ibrahim, Song et al., 2012,Guo, Chatterjee, Li et al., 2012,Turek, Joshu, Kohsaka et al., 2005). PER2 can directly repress PPARγ to inhibit adipocyte development (15). Notably, activation of the clock repressor Rev-erbα by specific agonists, a CLOCK/Bmal1 direct target gene, demonstrated anti-obesity efficacy in mice with improvement of dyslipidemia (16). On the other hand, RORα modulation by a natural ligand, Nobiletin, inhibited adipogenesis with in vivo effects on countering obesity (17). Thus, targeting the circadian clock function in modulating adipocyte development may provide novel pharmacological interventions for obesity and related metabolic consequences.

Through orchestration of time-of-day dependence of key metabolic pathways in distinct metabolic organs, circadian temporal control is intimately linked with metabolic homeostasis (2). Circadian misalignment predisposes to obesity and insulin resistance, as established by large-scale epidemiological investigations and accumulating experimental evidence (4, 18-22). Significant dampening of clock oscillation amplitude occurs with nutritional overload in high-fat diet-induced obesity, and clock disruption could be synergistic in inducing diabetes with over-nutrition (23, 24). It is thus conceivable that pharmacological targeting of key clock regulators to maintain or re-enforce clock-controlled metabolic rhythms may provide new avenues for anti-obesity therapies.

Through a high throughput screening pipeline, we recently discovered a new clock activator, chlorhexidine (CHX), and demonstrated its pro-myogenic efficacy (25). Here we show that, in distinct adipocyte progenitor models, CHX displayed strong action on inhibiting adipocyte differentiation. Via structural optimization, we obtained a CHX analog CM002 that activated clock with improved anti-adipogenic effects. Collectively, our findings revealed the potential for developing clock-activating compounds for anti-obesity interventions.

## Materials & Methods

### Animal studies

Mice were maintained in the City of Hope vivarium under a constant 12:12 light dark cycle provided with nestlets and maze for cage enrichment. All animal experiments were approved by the Institutional Animal Care & Use Committee (IACUC) of City of Hope and performed according to the IACUC approval. Male Bmal1-null mice in C57BL/6 background were obtained from the Jackson Laboratory (Strain #:009100). Mice were maintained in the lab and used for primary preadipocyte isolation at 10-12 weeks of age.

### Cell culture and adipogenic differentiation

3T3-L1 and C3H10T1/2 cell lines were obtained from ATCC and maintained in DMEM with 10% fetal bovine serum supplemented with 1% Penicillin-Streptomycin-Glutamine, as previously described (26, 27). 0.25% Trypsin was used for digestion and subculture of these cell lines. For adipogenic differentiation, induction media containing 1.6μM insulin, 1μM dexamethasone, 0.5mM IBMX and 0.5 uM Rosiglitazone was used for 3 days followed by maintenance medium with insulin for 3 days for 3T3-L1, and for 5 days for C3H10T1/2 cells, as previously described (17, 27).

### Generation of stable adipogenic progenitor cell lines containing Per2::dLuc

3T3-L1 preadipocytes and C3H10T1/2 mesenchymal stem cells obtained from ATCC were used for *Per2::dLuc* lentiviral transduction and stable clone selection using puromycin, as described previously (17, 28). Briefly, cells were transfected with lentiviral packaging plasmids (pSPAX.2 and pMD2.G) and lentivirus vectors *Per2::dLuc* using PEI Max (Polysciences). At 48 hours post-transfection, lentiviruses were collected. 3T3-L1 and C3H10T1/2 U2OS cells were infected using collected lentiviral media supplemented with polybrene. 24 hours following lentiviral infection, stable cell lines were selected in the presence of 2 μg/ml puromycin.

### Primary preadipocyte isolation and adipogenic induction

The stromal vascular fraction containing preadipocytes were isolated from subcutaneous fat pads, as previously described (29). Briefly, fat pads were cut into small pieces and digested using 0.1% collagenase Type 1 with 0.8% BSA at 37^0^C with constant shaking for 60 minutes. The digested homogenate was passed through Nylon mesh and centrifuged to collect the pellet containing the stromal vascular fraction with preadipocytes. Pelleted preadipocytes were cultured in F12/DMEM supplemented with bFGF (2.5 ng/ml), expanded for two passages and subjected to differentiation in 6-well plates at 90% confluency. Adipogenic differentiation was induced for 2 days in medium containing 10% FBS, 1.6 μM insulin, 1 μM dexamethasone, 0.5 mM IBMX, 0.5 uM rosiglitazone before switching to maintenance medium for 4 days with insulin only. Compounds at indicated concentrations were administered for the entire differentiation time course following adipogenic induction.

### Oil-red-O and Bodipy staining

Stainings for neutral lipids during adipogenic differentiation were performed as previously described (26). Briefly, for oil-red-O staining, cells were fixed with 10% formalin and incubated in 0.5% oil-red-O solution for 1 hour. Bodipy 493/503 was used at 1mg/L together with DAPI for 15 minutes, following 4% paraformaldehyde fixation and permeabilization with 0.2% triton-X100.

### Continuous Bioluminescence monitoring of Per2::dLuc luciferase reporter

3T3-L1, C3H10T1/2 or primary preadipocytes containing a *Per2::dLuc* luciferase reporter were used for bioluminescence recording, as previously described (17). Cells were seeded at 4x10^5^ density on 24 well plates at 90% confluence following overnight culture with explant medium luciferase recording media. Explant medium contains DMEM buffer stock, 10% FBS, 1% PSG, pH7 1M HEPES, 7.5% Sodium Bicarbonate, Sodium Hydroxide (100mM) and XenoLight D-Luciferin bioluminescent substrate (100mM). Raw and subtracted results of real-time bioluminescence recording data for 6 days were exported, and data was calculated as luminescence counts per second, as previously described (30). LumiCycle Analysis Program (Actimetrics) was used to determine clock oscillation period, length amplitude and phase. Briefly, raw data following the first cycle from day 2 to day 5 were fitted to a linear baseline, and the baseline-subtracted data (polynomial number = 1) were fitted to a sine wave, from which period length and goodness of fit and damping constant were determined. For samples that showed persistent rhythms, goodness-of-fit of >80% was usually achieved.

### TOPFlash luciferase reporter assay

M50 Super 8xTOPFlash luciferase reporter containing Wnt-responsive TCF bindings sites was a gift from Randall Moon (31) provided by Addgene (Addgene plasmid # 12456). For transient transfection with luciferase reporter, cells were seeded and grown overnight to reach 90% confluency. 24 hours following transfection, 10% Wnt3a conditioned media obtained from L Wnt-3A cell line (ATCC CRL-2647) was added to induce Wnt signaling. Luciferase activity was assayed using Dual-Luciferase Reporter Assay Kit (Promega) in 96-well black plates. TOPFlash luciferase reporter luminescence was measured on microplate reader (TECAN infinite M200pro) and normalized to control FOPFlash activity, as previously described (28). The mean and standard deviation values were calculated for each well using four replicates and graphed.

### Immunoblot analysis

Total protein was extracted using lysis buffer containing 3% NaCl, 5% Tris-HCl, 10% Glycerol, 0.5% Triton X-10 in MilliQ water with protease inhibitor cocktail. 20-40 µg of total protein was resolved on 10% SDS-PAGE gels followed by immunoblotting on PVDF membranes (Bio-rad). Membranes were developed by chemiluminescence (SuperSignal West Pico, Pierce Biotechnology) and signals were obtained via a chemiluminescence imager (Amersham Imager 680, GE Biosciences). Primary antibodies used are listed in Supplemental Table 1.

### RNA extraction and RT-qPCR analysis

PureLink RNA Mini Kit (Invitrogen) were used to isolate total RNA from cells. cDNA was generated using Revert Aid RT kit (ThermoFisher) and quantitative PCR was performed using SYBR Green Master Mix (Thermo Fisher) in triplicates on ViiA 7 Real-Time PCR System (Applied Biosystems). Relative gene expression was calculated using the comparative Ct method with normalization to 36B4 as internal control. PCR primers sequence are listed in Supplemental Table 2.

***Statistical analysis*** Data are presented as mean ± SD. Each experiment was repeated at minimum three times to validate the result. The number of replicates were indicated for each experiment in figure legends. Two-tailed Student’s t-test or One-way ANOVA with post-hoc analysis for multiple comparisons were performed as appropriate as indicated using GraphPad PRISM. P<0.05 was considered statistically significant.

## Results

### Chlorhexidine activation of adipocyte-intrinsic circadian clock

Circadian clock exerts coordinated control in adipogenesis, and Bmal1, through its transcriptional control of the Wnt signaling pathway, inhibits adipocyte development(28, 32). Based on our recent study that identified CHX as a novel clock-activating molecule by promoting CLOCK/Bmal1 transcriptional activity (25), we tested whether it could modulate cell-intrinsic clock function in adipogenic precursor cells. Using the mesenchymal precursor C3H10T1/2 (10T1/2) and 3T3-L1 preadipocytes as adipogenic progenitor models, we generated stable cell lines containing the *Per2::dLuc* luciferase reporter to examine CHX effect on clock properties. In 10T1/2 mesenchymal precursor cells, CHX treatment at 1 and 2 μM led to a dose-dependent shortening of clock period length (Fig. 1A& 1B). In addition, CHX at these concentrations were able to augment clock cycling amplitude (Fig. 1C), consistent with its function as a clock activator. The effects of CHX on reducing period length and increasing amplitude were also observed in lineage-committed 3T3-L1 preadipocytes at 1 μM (Fig. S1). Using primary preadipocytes isolated from the *Per2:dLuc* knock-in mice (33), we further examined CHX modulation of clock properties. CHX exhibited a similar effect on shortening period length (Fig. 1D & 1E), although it failed to modulate amplitude in this primary adipogenic precursor cell type (Fig. 1F). Interestingly, analysis of clock gene regulations by CHX in 10T1/2 cells revealed inductions of *Bmal1* together with CLOCK/Bmal1 direct target genes, *Nr1d1*, *Cry2* and *Per2*, in line with activation of CLOCK/Bmal1-mediated transcription (Fig. 1G). Largely similar effects on inducing core clock genes were observed in 3T3-L1 adipocytes (Fig. 1H).

**Figure 1.**
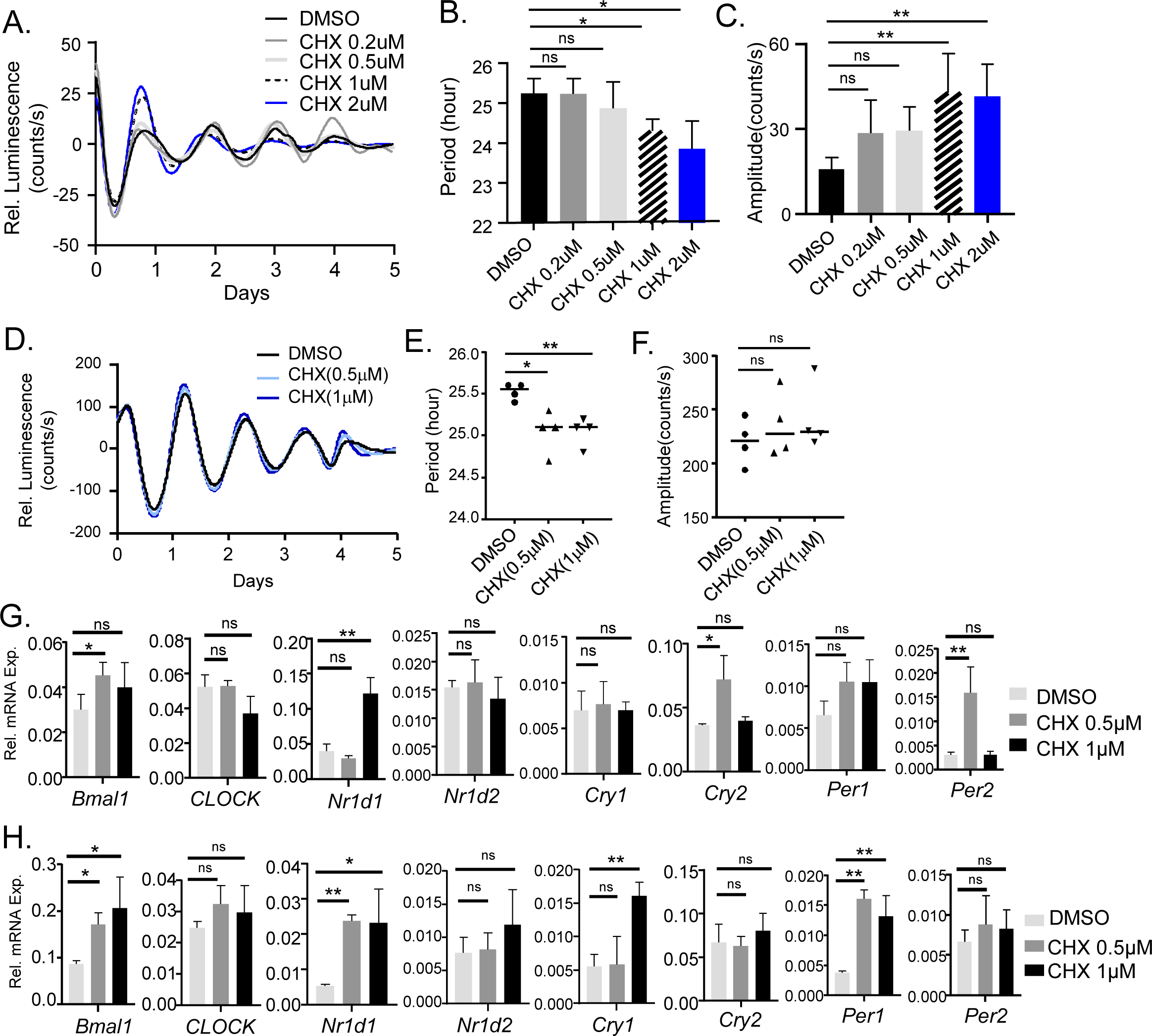
Chlorhexidine activity as a clock activator in adipogenic precursor cells. (A-F) Baseline-adjusted tracing plots of average luciferase bioluminescence activity of *Per2::dLuc* reporter-containing C3H10T1/2 (A), and primary preadipocytes isolated from *Per2::dLuc* knock-in mice (D) for 5 days, with corresponding quantitative analysis of clock period length (B, E) and cycling amplitude (C, F). Chlorhexidine (CHX) were added at indicated concentrations at the start of luciferase recording. Data are presented as Mean ± SD of n=4 replicates for each concentration tested. (G, H) RT-qPCR analysis of clock gene expression at indicated concentrations of CHX treatment for 6 hours in C3H10T/2 (G), and 3T3-L1 preadipocytes (H). Data are presented as Mean ± SD of n=3 replicates. *, **: p<0.05 and 0.01 CHX vs. DMSO by Student’s t test.

### Chlorhexidine inhibits differentiation of adipogenic mesenchymal progenitors

Based on previous findings of circadian clock regulation of adipogenesis, we tested whether CHX, as a clock activator, impacts adipocyte development. During induction of adipogenesis in 10T1/2 mesenchymal precursors, CHX treatment led to a dose-dependent inhibition of adipocyte maturation at the early stage of differentiation at 5 days after adipogenic induction, as shown by phase-contrast, oil-red-O and Bodipy staining of lipid accumulation as a marker for mature adipocyte formation (Fig. 2A & 2B). This inhibitory effect of CHX on 10T1/2 adipogenic differentiation remained evident at later stages as indicated by oil-red-O (Fig. 2C) and Bodipy staining (Fig. 2D & 2E). Further analysis of the adipogenic program demonstrated significantly impaired levels of adipogenic factors in day 8-differentiated C3H10T1/2 cells, including CEBP/α and PPARγ (Fig. 2F & 2G), with attenuated expression of fatty acid synthase (FASN). These effects of CHX revealed its suppression of the lineage commitment and maturation of adipogenic mesenchymal precursor cells.

**Figure 2.**
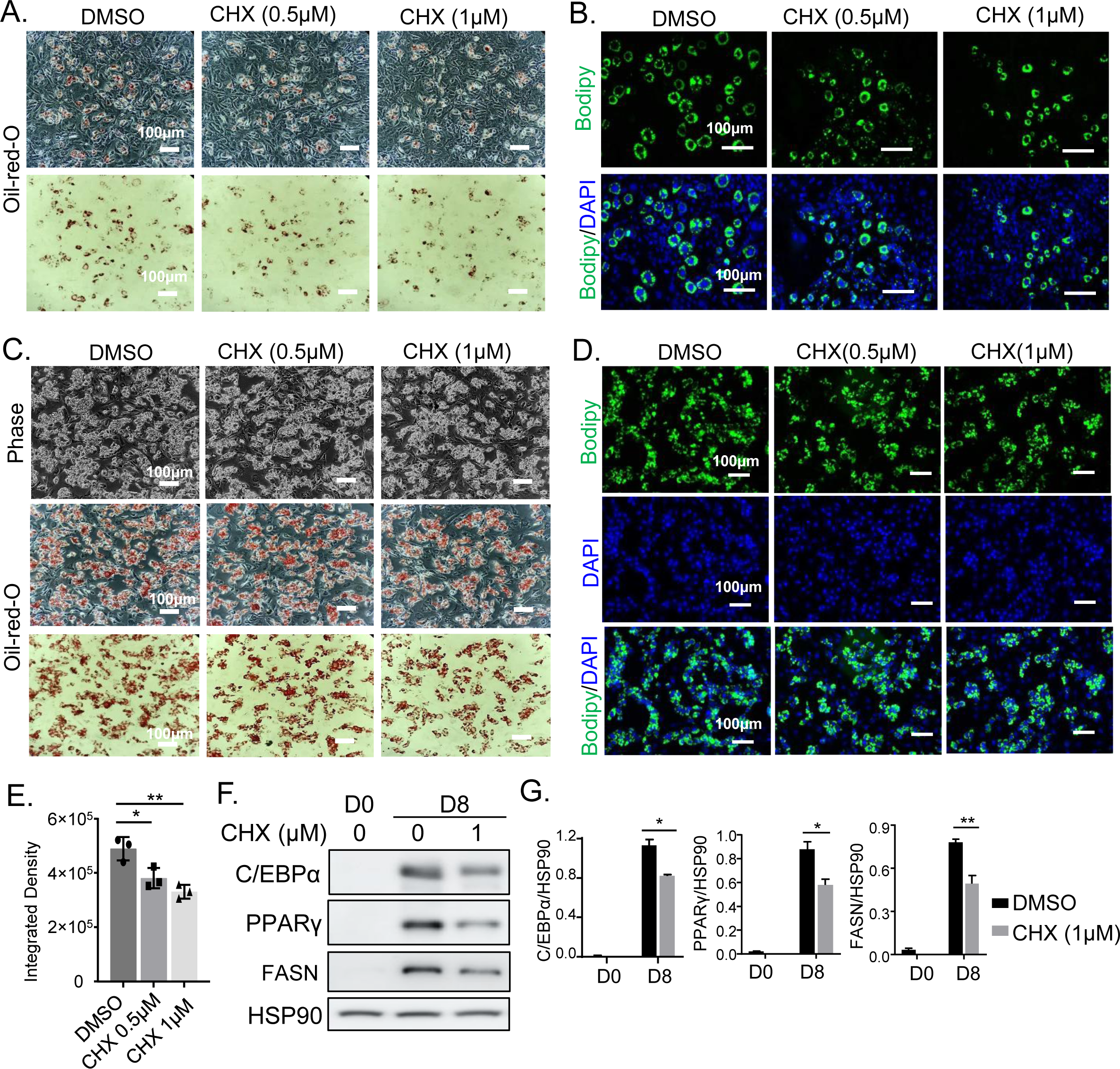
Chlorhexidine inhibition of lineage commitment and differentiation of adipogenic mesenchymal progenitors. (A, B) Representative images of oil-red-O staining (A) and Bodipy staining (B) of C3H10T1/2 cells at 5 days following adipogenic induction, treated with or without chlorhexidine at indicated concentrations. (C, D) Representative images of phase-contrast and oil- red-O staining (C) and Bodipy staining (D) of C3H 10T1/2 cells at 8 days following adipogenic induction, treated with or without Chlorhexidine at indicated concentrations. (E) Average intensity of BODIPY fluorescence of three representative fields as shown in D was obtained using Image J and normalized to DAPI signal. Scale bars: 100 μm. (F, G) Representative images of immunoblot analysis of adipogenic program before and after 7 days of 10T1/2 adipogenic differentiation with or without 1μM of Chlorhexidine (F), with quantification normalized to HSP90 level (G). Each lane represents three pooled samples. *, **: p<0.05 and 0.01 CHX vs. DMSO by Student’s t test.

### Chlorhexidine inhibition of preadipocyte terminal differentiation

We next determined whether CHX inhibits terminal differentiation of lineage-committed 3T3-L1 preadipocytes. In these adipogenic progenitors, CHX was sufficient to reduce the formation of lipid-laden mature adipocytes in a dose-dependent manner, as indicated by phase-contrast and oil-red-O staining following 6 days of adipogenic induction (Fig. 3A). A similar result of CHX inhibitory effect on adipocyte development was demonstrated via Bodipy staining (Fig. 3B & 3C). Interestingly, at day 6 of differentiation, examination of adipogenic factor induction revealed only moderate changes toward lower C/EBPα and PPARγ protein levels in cells treated with CHX that were not statistically significant (Fig. 3D & 3E). We next determined CHX effect on terminal differentiation using primary preadipocytes isolated from the stromal vascular fraction of adipose depot. Following six days of differentiation, CHX at 0.5 and 1 µM displayed a marked dose-dependent, effect on suppressing mature adipocyte formation, as revealed by phase-contrast morphology, oil-red-O staining (Fig. 4A & Fig. S3A), or Bodipy staining (Fig. 4B, 4C and S3B). Similar to the observation from 3T3-L1 preadipocytes, CHX only resulted in a tendency toward suppressing adipogenic factor induction at day 6 of differentiation (Fig. 4D & 4E). However, CHX markedly attenuated the terminal differentiation marker of adipocytes, FASN, suggesting its modulation of terminal differentiation without significantly altering the adipogenic program. Given the effect of CHX on attenuating differentiation of these lineage-committed preadipocytes, 3T3-L1 and primary preadipocytes, it may modulate adipogenesis by influencing early lineage determination as well as maturation of adipogenic progenitors.

**Figure 3.**
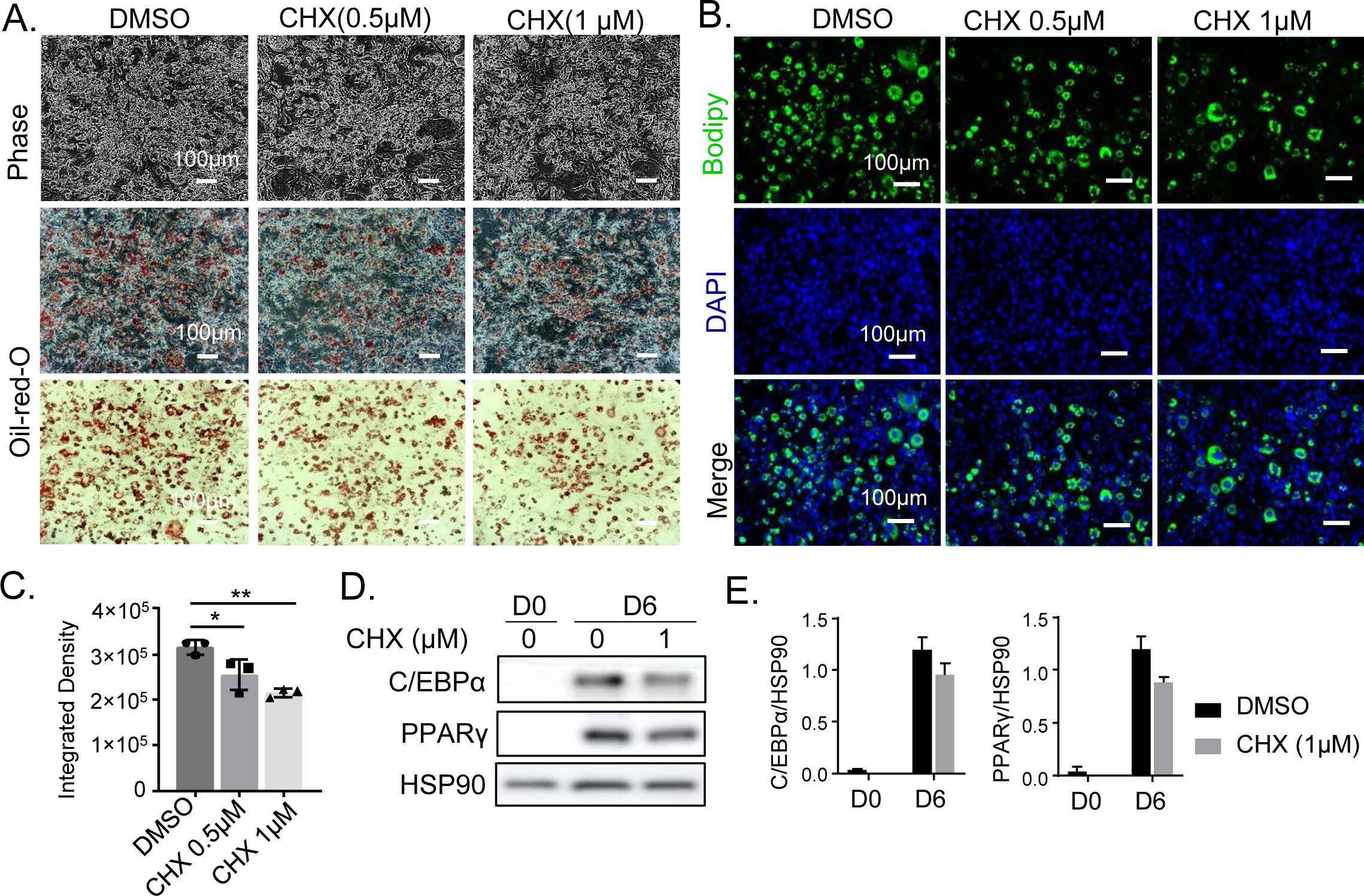
Chlorhexidine inhibitory effect on terminal differentiation of preadipocytes. (A-C) Representative images of phase-contrast and oil-red-O staining (A), and Bodipy fluorescence staining (B) with quantitative analysis (C), of adipogenic differentiation of 3T3-L1 preadipocytes at 6 days of differentiation treated with or without CHX at indicated concentrations. Scale bar: 100 μm. *, **: p<0.05 and 0.01 CHX vs. DMSO by Student’s t test. (D, E) Representative images of immunoblot analysis of adipogenic gene expression before (day 0) and after day 6 of differentiation (D) with quantification normalized to HSP90 level (E).

**Figure 4.**
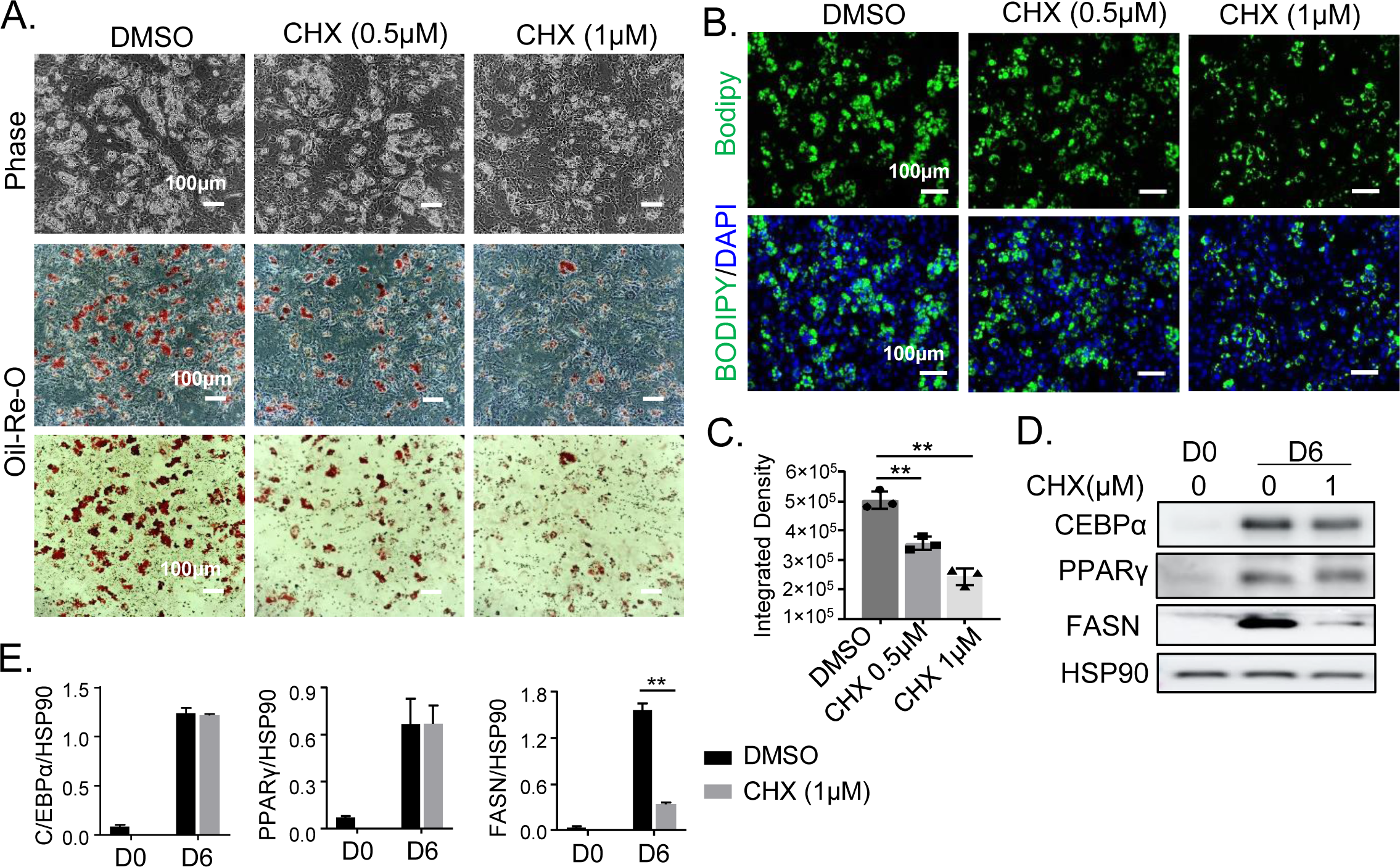
Effect of Chlorhexidine on suppressing primary preadipocyte differentiation. (A- C) Representative images of phase-contrast and oil-red-O staining (A) and Bodipy fluorescence staining (B) with quantitative analysis (C) of primary preadipocytes at day 6 of adipogenic differentiation treated with CHX at indicated concentrations. Scale bar: 100 μm. (D, E) Representative images of immunoblot analysis of adipogenic gene expression before (day 0) and after 6 days of primary adipocyte differentiation (D), with quantitative analysis normalized to HSP90 level (E). **: p< 0.01 CHX vs. DMSO by Student’s t test.

### Chlorhexidine induction of Wnt signaling pathway in adipogenic precursors

Wnt signaling pathway is a potent inhibitory development signal for adipose tissue development and adipogenic differentiation (34). Based on prior studies of Bmal1 transcriptional control of the Wnt pathway, we postulated that CHX activation of CLOCK/Bmal1 target genes in Wnt signaling may underlie its negative modulation of adipogenesis. Through RT-qPCR analysis, we examined Wnt signaling components that are direct targets of Bmal1 we previously identified (28, 29). CHX induced the mRNA levels of *Wnt1*, *Wnt10a*, *Frizzled 2* (*Fzd2)*, *Frizzled 5* (*Fzd5)* that are up-stream Wnt ligands and receptors in 10T1/2 cells (Fig. 5A). In 3T3-L1 preadipocytes, similar regulations of Wnt ligands and *Fzd5* were observed, together with inductions of *Disheveled 2* (*Dvl2*) and *β-catenin* expression (Fig. 5B). Consistent with its modulation of β-catenin transcript, CHX treatment in 10T/2 cells was sufficient to re-activate the loss of β-catenin protein level during adipogenic differentiation (Fig. 5C & 5D). Furthermore, using a Wnt-responsive TOPFlash luciferase reporter containing TCF4 bindings sites to assay for Wnt signaling activity, we found that CHX at 0.2 and 0.5 µM induced the luciferase reporter activity ∼3 fold at either basal or Wnt-stimulated conditions (Fig. 5E), suggesting CHX activation of Wnt signaling in adipogenic progenitors that suppresses differentiation.

**Figure 5.**
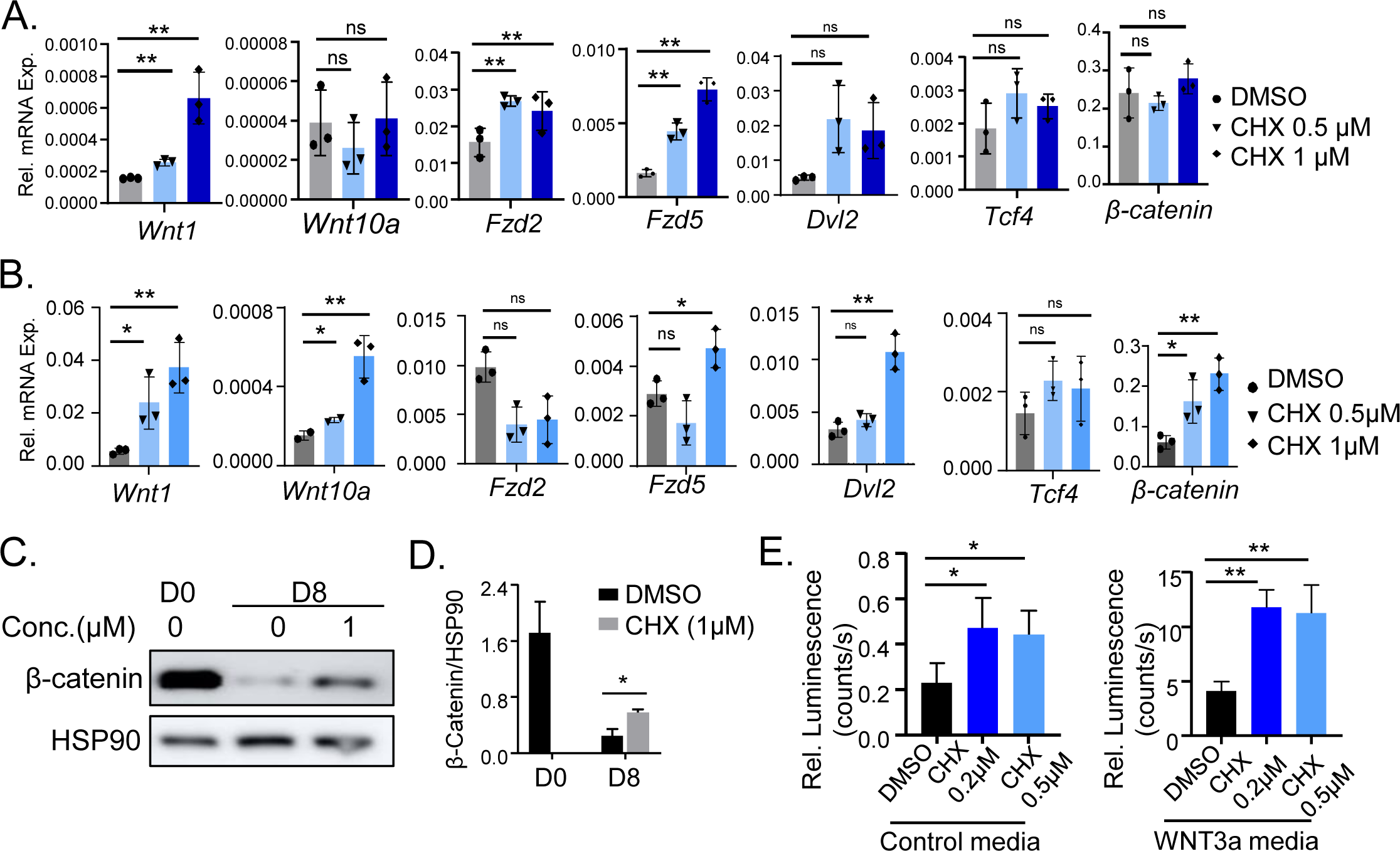
Chlorhexidine promotes Wnt signaling in adipogenic progenitors. (A, B) RT-qPCR analysis of CHX effect on expression of key components of Wnt signaling pathway in C3H10T1/2 cells (A), and 3T3-L1 preadipocytes (B), treated with indicated concentrations of chlorhexidine for 6 hours. Data are presented as Mean ± SD of n=3 replicates. *, **: p<0.05 and 0.01 CHX vs. DMSO by Student’s t test. (C, D) Representative images of immunoblot analysis of CHX effect (1µM) on β-catenin protein level in C3H10T1/2 cells before and after adipogenic induction for 8 days (C), with quantitative analysis normalized to HSP90 level (D). (E) Luciferase reporter assay of Wnt signaling pathway activity by a Wnt-responsive TOPFlash luciferase reporter treated with control or Wnt3a-containing media at 10% concentration. Data are presented as Mean ± SD of n=4 replicates. *, **: p<0.05 or 0.01 CHX vs. DMSO by Student’s t test.

### Chlorhexidine structural analog CM002 displays clock-activating property

Based on the chemical scaffold of Chlorhexidine, we made modification of specific chemical groups to generate structural analogs via chemical synthesis. Using an *Per2::dLuc* luciferase-containing U2OS reporter cell line, we screened the clock-modulatory activities of these compounds that led to identification of CM002, a molecule with approximately half of the structural scaffold of CHX (Fig. S2A), as a new clock activator. CHX was originally identified via a high through-put screening for clock modulators against a hydrophobic pocket of the CLOCK protein (25). Through molecular docking analysis, it revealed potential CM002 occupation within the shared hydrophobic pocket of CLOCK with CHX (Fig. 6A & 6B). Consistent with the overlap of CM002 structure with CHX scaffold, its predicted binding surface coincides with a large portion of CHX interactions within the CLOCK protein (Fig. S2B). A detailed examination of CLOCK protein residues with potential interactions with CM002 within 3-4A^0^ were illustrated (Fig. 6B). As shown using the U2OS *Per2::dLuc* reporter cells, CM002 exhibited a dose-dependent effect on inducing clock period length shortening (Fig. 6C & 6D) without significantly affecting cycling amplitude (Fig. 6E), demonstrating its clock-activating property. Furthermore, CM002 treatment of 10T1/2 cells led to induction of core clock genes, *CLOCK* and *Bmal1,* together with up- regulation of its direct target within the molecular clock loop, *Nr1d1* (Fig. 6F). Given that clock exerts transcriptional control of Wnt pathway and CHX can induce Wnt signaling due to clock activation, we examined whether CM002 modulates Wnt activity using the TOPFlash reporter (31). Under basal condition, CM002 treatment at 0.5 µM was sufficient to stimulate reporter activity to a similar degree as CHX (Fig. 6G). Upon Wnt3a media stimulation, CM002 at lower concentrations of 0.1 and 0.2 µM also induced TOPFlash activity comparable to that of CHX at 0.5 µM, suggesting improved Wnt-activating efficacy of CM002 (Fig. 6H).

**Figure 6.**
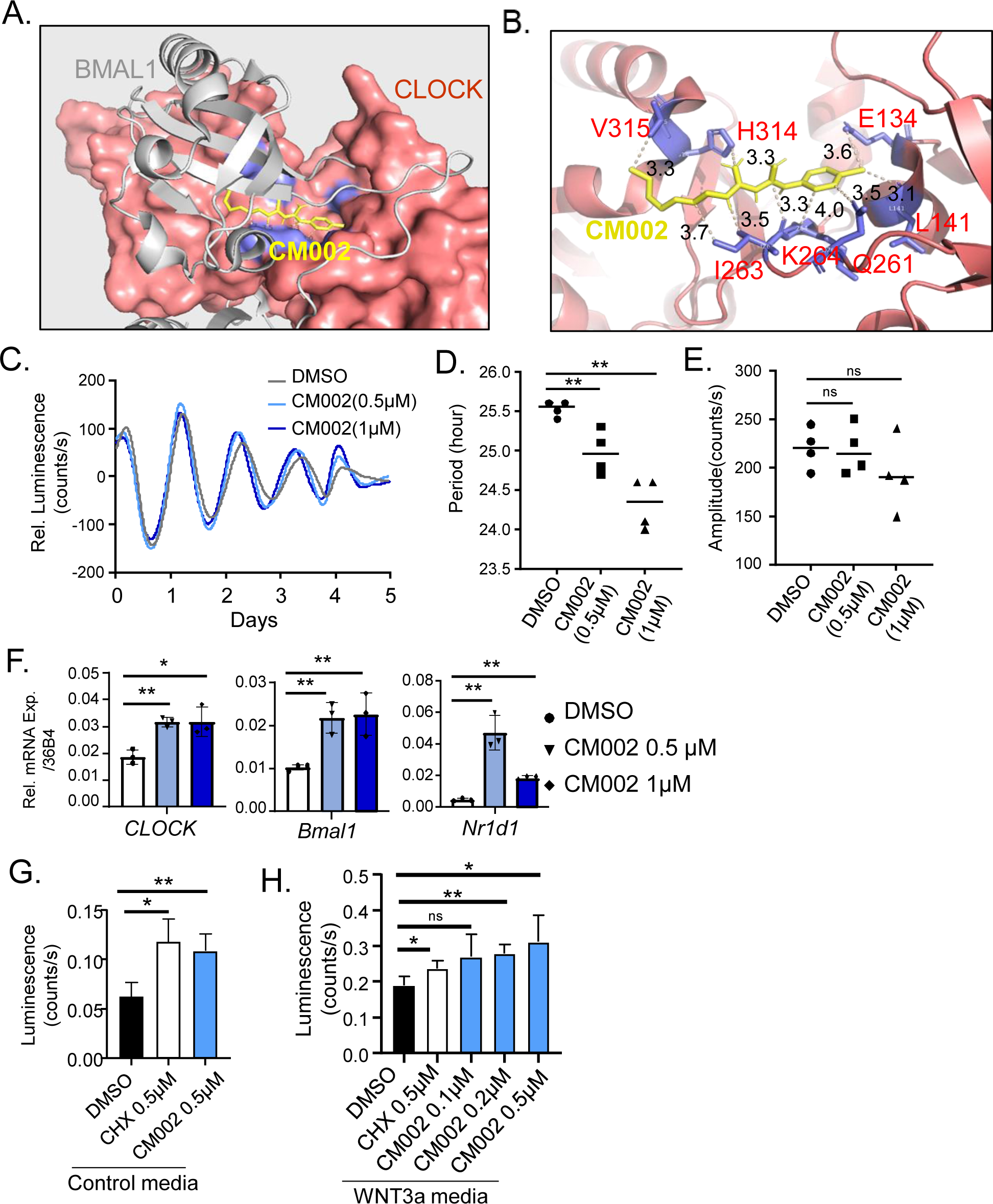
CM002 as a chlorhexidine analog with clock-activating properties. (A) Docking conformation of CM002 within the CLOCK protein hydrophobic pocket. Crystal structure shown for CLOCK (red, surface mode) with Bmal1 (grey, cartoon mode) is based on PDB: 4f3l. (B) Predicted CM002 interactions with CLOCK protein residues within 3-4A^0^ distance. (C-E) Baseline-adjusted tracing plots of average bioluminescence activity of U2OS cell line with stable expression of a *Per2::dLuc* reporter for 5 days (C), with quantitative analysis of clock period length (D) and cycling amplitude (E). CM002 at indicated concentrations were added at start of luciferase recording. Data are presented as Mean ± SD of n=4 replicates for each concentration tested. (F) RT-qPCR analysis of clock gene expression of C3H10T/2 cells treated by CM002 at indicated concentrations for 6 hours. Data are presented as Mean ± SD of n=3 replicates. *, **: p<0.05 and 0.01 CHX vs. DMSO by Student’s t test. (G, H) Analysis of CM002 effect on Wnt signaling activity using TOPFlash luciferase reporter activity assay under basal media (G) or Wnt3a- stimulated condition (H). Data are presented as Mean ± SD of n=4 replicates. *, **: p<0.05 or 0.01 CM002 vs. DMSO by Student’s t test.

### CM002 inhibits adipogenesis of mesenchymal precursor and committed preadipocytes

As a CHX analog that activated clock and stimulated Wnt signaling, we postulated that CM002 may have anti-adipogenic effects as shown for CHX. Indeed, in 10T1/2 adipogenic precursor cells, CM002 markedly suppressed their differentiation in a dose-dependent manner as shown by oil-red-O (Fig. 7A) or Bodipy staining of lipids (Fig. 7B & 7C). Analysis of adipogenic induction at day 8 of differentiation revealed marked inhibitions of adipogenic factors C/EBPα and PPARγ, suggesting a robust block of differentiation (Fig. 7D & 7E). Protein levels of mature adipocyte markers, including FASN and fatty acid binding protein 4 (FABP4), were substantially reduced in CM002-treated cells as compared to controls, indicating impaired adipocyte maturation.

**Figure 7.**
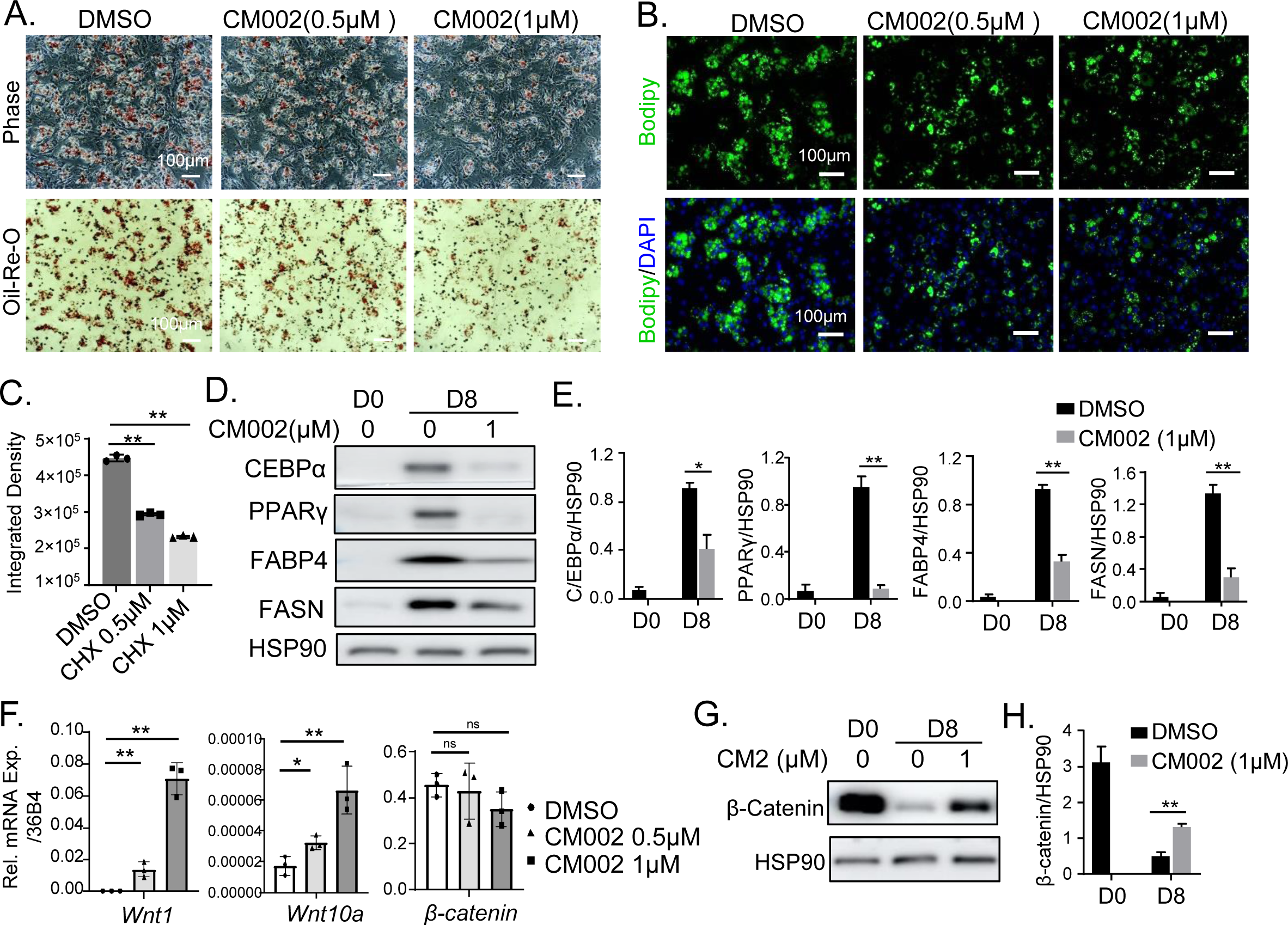
CM002 inhibits C3H10T1/2 mesenchymal adipogenic progenitor differentiation. (A-C) Representative images of oil-red-O (A), and Bodipy staining (B) with quantitative analysis (C) of 10T1/2 cells after 8 days of adipogenic differentiation treated with CM002 at indicated concentrations. Scale bar: 100 μm. (D, E) Representative images of immunoblot analysis of adipogenic program before and at day 8 of C3H10T1/2 differentiation treated with or without 1μM of CM002 (D), with quantification using normalization to HSP90 (E). Each lane represents three pooled samples. (F) RT-qPCR analysis of clock gene expression of C3H10T/2 cells treated with CM002 at indicated concentrations for 6 hours. N=3 replicates. (G, H) Representative immunoblot images of total β-catenin level before and at day 8 of C3H10T1/2 differentiation treated with 1μM of CM002 (G), with quantification using normalization to HSP90 (H).

Surveying of key Wnt signaling components revealed significant up-regulations of Wnt ligands including *Wnt1* and *Wnt10a*, although β-catenin was not altered (Fig. 7F). With adipogenic induction in these cells, β-catenin protein level demonstrated a marked decline as compared to undifferentiated state, as expected (Fig. 7G & 7H). Notably, CM002 partially reversed the loss of β-catenin protein expression at the differentiated state, suggesting its role in mediating CM002 inhibition of adipogenesis.

To determine the effect of CM002 on primary adipocyte differentiation and whether its anti-adipogenic activity is dependent on clock modulation, we used primary preadipocytes isolated from normal controls (BMCtr) and clock-deficient *Bmal1*-null (BMKO) mice. The administration of CM002 to differentiating primary preadipocytes from normal controls strongly inhibited their maturation as shown by oil-red-O staining (Fig. 8A). In *Bmal1*-null preadipocytes derived from BMKO mice, CM002 effect on reducing adipocyte formation was blunted. Bodipy staining further confirmed this finding, with dose-dependent reductions by CM002 at 0.5 and 1μM concentration, while this inhibition of adipocyte maturation was attenuated in BMKO preadipocytes (Fig. 8B & 8C). Thus, the effect of CM002 on inhibiting adipogenesis was mediated, at least in part, by a clock-dependent mechanism.

**Figure 8.**
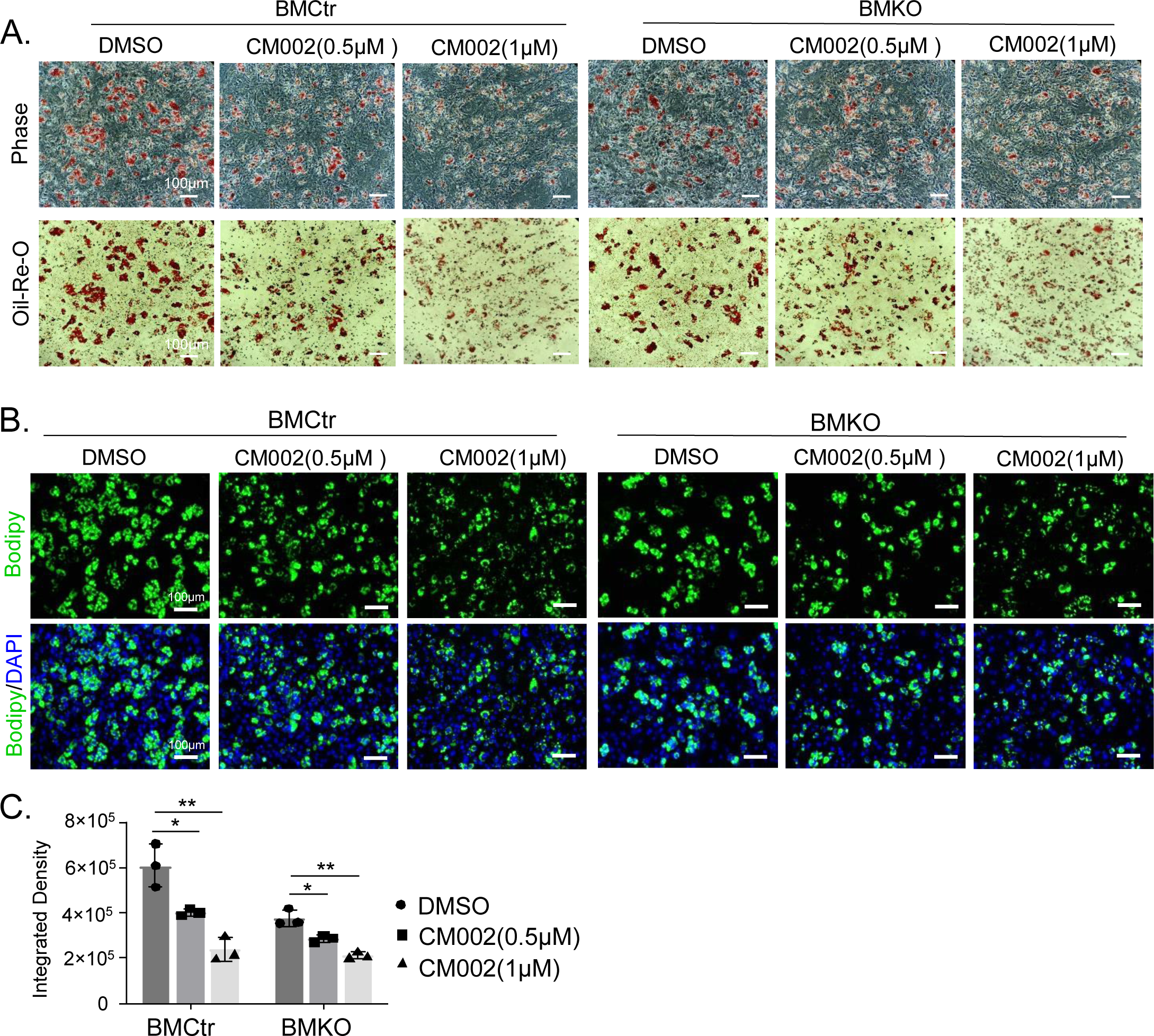
Effect of CM002 on inhibiting terminal differentiation of primary preadipocyte. (A-C) Representative images of oil-red-O (A), and Bodipy fluorescence staining (B) with quantitative analysis (C), of primary preadipocytes from control (BMCtr) or Bmal1-null mice (BMKO) after 6 days of adipogenic differentiation. Cells were treated with or without CM002 at indicated concentrations. Scale bars: 100 μm. *, **: p<0.05 or 0.01 CM002 vs. DMSO by Student’s t test.

## Discussion

Circadian clock plays a key regulatory role in modulating adipogenesis, and a large body of evidence indicates that clock disruption leads to the development of obesity and insulin resistance (3). In light of the widespread “social jetlag” in a modern lifestyle, circadian clock could be a potential therapeutic target to counter the current epidemic of metabolic diseases (7, 12). Employing distinct adipogenic progenitor cellular models, the current study uncovered the effect of a new clock-activating compound, chlorhexidine, in inhibiting adipocyte development, and further demonstrated the anti-adipogenic efficacy of a new analog.

Accumulating studies to date have established the intimate association between circadian clock disruption and the development of metabolic diseases (3, 35). Given the pervasive temporal control of circadian clock of rate-limiting enzymes of metabolic pathways and circadian oscillation of metabolites, dysregulation of various clock-controlled metabolic processes contributes to altered metabolic homeostasis. We previously reported that clock transcription factor Bmal1 exert direct transcriptional control of Wnt signaling pathway components to modulate its activity, a known developmental signal in suppressing adipogenesis (28). Loss of clock function, due to deficiency of the core clock transcription activators CLOCK or Bmal1 in mice, leads to obesity (28, 36, 37). In addition, disruption of clock via various environmental clock manipulations predispose to the development of obesity (19, 38-40), while shiftwork is linked with a strong risk for insulin resistance and type II diabetes (4, 20, 40). Thus targeting the specific link of clock with adipogenesis, particularly by maintaining or re-enforcing proper circadian clock control in the face of wide-spread circadian misalignment, may lead to new avenues for obesity treatment through clock modulation. Nobiletin, a clock amplitude-enhancing molecule by activating the positive regulator of the clock RORa (11), was found to suppress adipogenesis and display strong anti-obesity effect in mice (17). Using a high through-put screening platform to identify novel clock modulators, we recently discovered Chlorhexidine as a clock-activating compound (25). In line with the role of clock function in promoting myogenesis via Wnt signaling, chlorhexidine demonstrated a robust pro-myogenic effect in both C2C12 and primary myoblasts with induction of Wnt activity. Importantly, the effect of chlorhexidine on promoting myogenic differentiation was mediated by clock, as this effect was completely abolished in *Bmal1*-deficient myoblasts (Kiperman et al., 2023).

Given the role of clock in promoting Wnt signaling to suppress adipogenesis (Guo et al., 2012), we tested chlorhexidine as a chemical probe to target clock function in adipocyte development. Current results revealed that pharmacological interventions to augment clock activity, via chlorhexidine or a new structural derivative, could suppress adipocyte maturation in multiple adipogenic progenitor populations tested. Consistent with its activity observed in myogenic progenitors, Chlorhexidine was able to activator clock function in these adipogenic progenitor cell types with stimulation of Wnt signaling. These findings suggest the potential for targeting clock function to inhibit adipogenic drive for anti-obesity effect, although rigorous in vivo studies are needed.

In an effort to optimize compound structure to achieve better efficacy and minimize toxicity, we generated new analogs through chemical modifications of the chlorhexidine scaffold and screened for molecules with improved clock-modulatory activity. This led to the identification of CM002 as a new clock activator. Clock activity monitoring revealed that CM002 displayed improved efficacy range on inducing clock period shortening without cellular toxicity within the concentrations tested, and its activity in promoting Wnt signaling was consistent with improved clock-modulatory efficacy. Notably, CM002 effect on adipogenic inhibition appears to be more robust than that of chlorhexidine, as shown by strong inhibition of the adipogenic gene program. More detailed comparisons between CM002 and CHX are needed in future studies to determine the relative anti-adipogenic efficacy, particularly in vivo. We observed certain cellular toxicity of CHX beyond 1 μM concentration, while the viability of cells treated by CM2 at this concentration was maintained (data not shown), suggesting potential wider concentration range for this compound. The precise effective range of these compounds and their safety profiles, potentially in preclinical models, requires future detailed investigations.

An intriguing finding from our study is the differential modulation of CHX and CM002 on adipogenic gene induction between the mesenchymal and lineage-committed progenitors, the C3H10T1/2 cells as compared to 3T3-L1 and primary preadipocytes. Both CHX and CM002 had robust effects on reducing adipocyte formation in mesenchymal progenitors, with consistent reduction of expression levels of key adipogenic factor and mature adipocyte markers. Interestingly, despite a comparable level of inhibition of adipocyte maturation by these molecules in 3T3-L1 and primary preadipocytes as indicated by the degree of lipid accumulation, their effects are mostly confined to blockade of mature adipocyte marker expression without significant modulation of adipogenic factors. These distinct regulations in specific progenitors suggest their potential developmental stage-specific modulation of the differentiation process. CHX and CM002 suppressed lineage commitment of adipogenic induction in mesenchymal progenitors with consequent impact on differentiation, whereas their effects on lineage-determined preadipocytes limited to terminal differentiation of adipocyte. These mechanisms of action of clock-activating molecules shed light on their therapeutic potentials to prevent preadipocyte conversion to mature adipocytes for anti-obesity drug development.

The anti-adipogenic effects of CHX and its derivatives may have anti-obesity potentials, although this requires to be further tested in vivo in appropriate pre-clinical model. It is also conceivable the further optimization of CHX or CM002 scaffolds are needed to obtain clock-activating compounds with anti-obesity efficacy in vivo. High-fat diet feeding-induced obesity leads to dampening of clock oscillatory amplitude, raising the possibility that promoting clock oscillation may reverse this effect of nutritional overload on clock-controlled metabolic pathways (24). Nobiletin, by activating the positive regulator of the clock RORa, was able to suppress adipogenesis with demonstrated anti-obesity effect in mice (17). Mediated by a common mechanism of transcriptional modulation of the Wnt signaling components, clock exerts anti-adipogenic effect while promotes myogenesis (25, 28, 29). This raises an intriguing possibility that maintaining or promoting proper clock modulation may have applications toward therapies against aging-associated sarcopenic obesity, characterized by the co-existing conditions of loss of skeletal muscle mass with adipose expansion in the elderly (41, 42). Altered clock function is associated with aging-related processes, with loss of cycling amplitude and a switch from rhythmic growth processes to skewed enrichment of stress response pathways in stem cell compartments (43, 44). Potential protective effects of augmenting clock regulation against aging-associated metabolic dysfunctions, or additional applications in aging-related pathologies, could be explored with discovery of diverse chemical modulators. Given the growing attention in targeting the circadian clock for disease treatment (45, 46), discovering clock activators to augment temporal control in specific biological processes could have therapeutic implications in a diverse array of human pathologies associated with circadian misalignment.

## Conclusions

In summary, our study uncovered, for the first time, the anti-adipogenic effects of new clock-modulating compounds that may have potential for anti-obesity drug development. With the widespread metabolic consequences of circadian clock disruption, pharmacological targeting of the clock machinery to maintain metabolic homeostasis may offer new avenues for the prevention or treatment of obesity and related metabolic consequences.

## Supporting information

SuppIementalFile

## List of Abbreviations

CLOCK: Circadian Locomotor Output Cycles Kaput
Bmal1: Brain and Muscle Arnt-like Protein 1
PAS: Per-Arnt-Sim
ROR: RAR-related Orphan Receptor
CHX: chlorhexidine
MyHC: myosin heavy chain

## Declarations

### Ethics approval and consent to participate

All animal experiments in this study were approved by the Institutional Animal Care & Use Committee (IACUC) of City of Hope according to the approval. The protocol number: 17110, entitled “circadian regulation of metabolism” with approval date is from 12/11/2020 to 12/10/2023.

### Consent for publication

Not applicable

### Availability of data and material

All data generated and analyzed during this study are included in this published article and associated Supplementary Information files.

### Authors’ contributions

XX, TK and WL: data curation and investigation, formal analysis, manuscript editing; ZF, AA WH and DH: data curation and manuscript editing; KM: formal analysis, project administration, manuscript writing and editing, and funding acquisition.

## Acknowledgements

We thank Drs. Steve Kay and Meng Qu at the University of Southern California for sharing luciferase reporter cell lines used in this study, and Drs. Seung-Hee Yoo and Zheng Chen at University of Texas at Houston Health Science Center for providing the Per2-luciferase plasmid.

## REFERENCE

1. Takahashi JS. Transcriptional architecture of the mammalian circadian clock. Nature reviews Genetics. 2017;18(3):164–79.

2. Bass J, Takahashi JS. Circadian integration of metabolism and energetics. Science (New York, NY. 2010;330(6009):1349-54.

3. Stenvers DJ, Scheer F, Schrauwen P, la Fleur SE, Kalsbeek A. Circadian clocks and insulin resistance. Nat Rev Endocrinol. 2019;15(2):75–89.

4. Pan A, Schernhammer ES, Sun Q, Hu FB. Rotating night shift work and risk of type 2 diabetes: two prospective cohort studies in women. PLoS medicine. 2011;8(12):e1001141.

5. Fonken LK, Workman JL, Walton JC, Weil ZM, Morris JS, Haim A, et al. Light at night increases body mass by shifting the time of food intake. Proceedings of the National Academy of Sciences of the United States of America. 2010;107(43):18664–9.

6. Sulli G, Lam MTY, Panda S. Interplay between Circadian Clock and Cancer: New Frontiers for Cancer Treatment. Trends Cancer. 2019;5(8):475–94.

7. Sulli G, Manoogian ENC, Taub PR, Panda S. Training the Circadian Clock, Clocking the Drugs, and Drugging the Clock to Prevent, Manage, and Treat Chronic Diseases. Trends Pharmacol Sci. 2018;39(9):812–27.

8. Cederroth CR, Albrecht U, Bass J, Brown SA, Dyhrfjeld-Johnsen J, Gachon F, et al. Medicine in the Fourth Dimension. Cell Metab. 2019;30(2):238–50.

9. Sulli G, Rommel A, Wang X, Kolar MJ, Puca F, Saghatelian A, et al. Pharmacological activation of REV-ERBs is lethal in cancer and oncogene-induced senescence. Nature. 2018;553(7688):351-5.

10. Oshima T, Niwa Y, Kuwata K, Srivastava A, Hyoda T, Tsuchiya Y, et al. Cell-based screen identifies a new potent and highly selective CK2 inhibitor for modulation of circadian rhythms and cancer cell growth. Sci Adv. 2019;5(1):eaau9060.

11. He B, Nohara K, Park N, Park YS, Guillory B, Zhao Z, et al. The Small Molecule Nobiletin Targets the Molecular Oscillator to Enhance Circadian Rhythms and Protect against Metabolic Syndrome. Cell metabolism. 2016;23(4):610–21.

12. Chen Z, Yoo SH, Takahashi JS. Development and Therapeutic Potential of Small-Molecule Modulators of Circadian Systems. Annu Rev Pharmacol Toxicol. 2018;58:231–52.

13. Stratmann M, Schibler U. REV-ERBs: more than the sum of the individual parts. Cell metabolism. 2012;15(6):791–3.

14. Preitner N, Damiola F, Lopez-Molina L, Zakany J, Duboule D, Albrecht U, et al. The orphan nuclear receptor REV-ERBalpha controls circadian transcription within the positive limb of the mammalian circadian oscillator. Cell. 2002;110(2):251–60.

15. Grimaldi B, Bellet MM, Katada S, Astarita G, Hirayama J, Amin RH, et al. PER2 controls lipid metabolism by direct regulation of PPARgamma. Cell metabolism. 2010;12(5):509–20.

16. Solt LA, Wang Y, Banerjee S, Hughes T, Kojetin DJ, Lundasen T, et al. Regulation of circadian behaviour and metabolism by synthetic REV-ERB agonists. Nature. 2012;485(7396):62- 8.

17. Xiong X, Kiperman T, Li W, Dhawan S, Lee J, Yechoor V, et al. The Clock-modulatory Activity of Nobiletin Suppresses Adipogenesis Via Wnt Signaling. Endocrinology. 2023;164(8).

18. McHill AW, Melanson EL, Higgins J, Connick E, Moehlman TM, Stothard ER, et al. Impact of circadian misalignment on energy metabolism during simulated nightshift work. Proceedings of the National Academy of Sciences of the United States of America. 2014;111(48):17302–7.

19. Scheer FA, Hilton MF, Mantzoros CS, Shea SA. Adverse metabolic and cardiovascular consequences of circadian misalignment. Proceedings of the National Academy of Sciences of the United States of America. 2009;106(11):4453–8.

20. van Amelsvoort LG, Schouten EG, Kok FJ. Duration of shiftwork related to body mass index and waist to hip ratio. International journal of obesity and related metabolic disorders : journal of the International Association for the Study of Obesity. 1999;23(9):973–8.

21. Shan Z, Li Y, Zong G, Guo Y, Li J, Manson JE, et al. Rotating night shift work and adherence to unhealthy lifestyle in predicting risk of type 2 diabetes: results from two large US cohorts of female nurses. BMJ. 2018;363:k4641.

22. Vetter C, Devore EE, Ramin CA, Speizer FE, Willett WC, Schernhammer ES. Mismatch of Sleep and Work Timing and Risk of Type 2 Diabetes. Diabetes care. 2015;38(9):1707–13.

23. Qian J, Yeh B, Rakshit K, Colwell CS, Matveyenko AV. Circadian Disruption and Diet-Induced Obesity Synergize to Promote Development of beta-Cell Failure and Diabetes in Male Rats. Endocrinology. 2015;156(12):4426–36.

24. Kohsaka A, Laposky AD, Ramsey KM, Estrada C, Joshu C, Kobayashi Y, et al. High-fat diet disrupts behavioral and molecular circadian rhythms in mice. Cell metabolism. 2007;6(5):414–21.

25. Kiperman T, Li W, Xiong X, Li H, Horne D, Ma K. Targeted screening and identification of chlorhexidine as a pro-myogenic circadian clock activator. Stem Cell Res Ther. 2023;14(1):190.

26. Nam D, Guo B, Chatterjee S, Chen MH, Nelson D, Yechoor VK, et al. The adipocyte clock controls brown adipogenesis through the TGF-beta and BMP signaling pathways. J Cell Sci. 2015;128(9):1835–47.

27. Liu R, Xiong X, Nam D, Yechoor V, Ma K. SRF-MRTF signaling suppresses brown adipocyte development by modulating TGF-beta/BMP pathway. Mol Cell Endocrinol. 2020;515:110920.

28. Guo B, Chatterjee S, Li L, Kim JM, Lee J, Yechoor VK, et al. The clock gene, brain and muscle Arnt-like 1, regulates adipogenesis via Wnt signaling pathway. FASEB journal : official publication of the Federation of American Societies for Experimental Biology. 2012;26(8):3453–63.

29. Chatterjee S, Nam D, Guo B, Kim JM, Winnier GE, Lee J, et al. Brain and muscle Arnt-like 1 is a key regulator of myogenesis. J Cell Sci. 2013;126(Pt 10):2213–24.

30. Xiong X, Li W, Nam J, Qu M, Kay SA, Ma K. The actin cytoskeleton-MRTF/SRF cascade transduces cellular physical niche cues to entrain the circadian clock. J Cell Sci. 2022;135(19).

31. Veeman MT, Slusarski DC, Kaykas A, Louie SH, Moon RT. Zebrafish prickle, a modulator of noncanonical Wnt/Fz signaling, regulates gastrulation movements. Curr Biol. 2003;13(8):680–5.

32. Nam D, Yechoor VK, Ma K. Molecular clock integration of brown adipose tissue formation and function. Adipocyte. 2016;5(2):243–50.

33. Yoo SH, Yamazaki S, Lowrey PL, Shimomura K, Ko CH, Buhr ED, et al. PERIOD2::LUCIFERASE real-time reporting of circadian dynamics reveals persistent circadian oscillations in mouse peripheral tissues. Proceedings of the National Academy of Sciences of the United States of America. 2004;101(15):5339–46.

34. Ross SE, Hemati N, Longo KA, Bennett CN, Lucas PC, Erickson RL, et al. Inhibition of adipogenesis by Wnt signaling. Science (New York, NY. 2000;289(5481):950-3.

35. Sinturel F, Petrenko V, Dibner C. Circadian Clocks Make Metabolism Run. J Mol Biol. 2020;432(12):3680–99.

36. Paschos GK, Ibrahim S, Song WL, Kunieda T, Grant G, Reyes TM, et al. Obesity in mice with adipocyte-specific deletion of clock component Arntl. Nat Med. 2012;18(12):1768–77.

37. Turek FW, Joshu C, Kohsaka A, Lin E, Ivanova G, McDearmon E, et al. Obesity and metabolic syndrome in circadian Clock mutant mice. Science (New York, NY. 2005;308(5724):1043-5.

38. Xiong X, Lin Y, Lee J, Paul A, Yechoor V, Figueiro M, et al. Chronic circadian shift leads to adipose tissue inflammation and fibrosis. Mol Cell Endocrinol. 2021;521:111110.

39. Kolbe I, Leinweber B, Brandenburger M, Oster H. Circadian clock network desynchrony promotes weight gain and alters glucose homeostasis in mice. Mol Metab. 2019;30:140–51.

40. Barclay JL, Husse J, Bode B, Naujokat N, Meyer-Kovac J, Schmid SM, et al. Circadian desynchrony promotes metabolic disruption in a mouse model of shiftwork. PLoS One. 2012;7(5):e37150.

41. Batsis JA, Villareal DT. Sarcopenic obesity in older adults: aetiology, epidemiology and treatment strategies. Nat Rev Endocrinol. 2018;14(9):513–37.

42. Cleasby ME, Jamieson PM, Atherton PJ. Insulin resistance and sarcopenia: mechanistic links between common co-morbidities. J Endocrinol. 2016;229(2):R67–81.

43. Solanas G, Peixoto FO, Perdiguero E, Jardi M, Ruiz-Bonilla V, Datta D, et al. Aged Stem Cells Reprogram Their Daily Rhythmic Functions to Adapt to Stress. Cell. 2017;170(4):678–92 e20.

44. Welz PS, Benitah SA. Molecular Connections Between Circadian Clocks and Aging. J Mol Biol. 2020;432(12):3661–79.

45. Manoogian ENC, Panda S. Circadian rhythms, time-restricted feeding, and healthy aging. Ageing Res Rev. 2017;39:59–67.

46. Chaix A, Zarrinpar A, Miu P, Panda S. Time-restricted feeding is a preventative and therapeutic intervention against diverse nutritional challenges. Cell metabolism. 2014;20(6):991–1005.

