## Supplementary material for "Anti-adipogenic properties of clock activator chlorhexidine and a new derivative": SuppIementalFile

### Supplemental Figure S1.

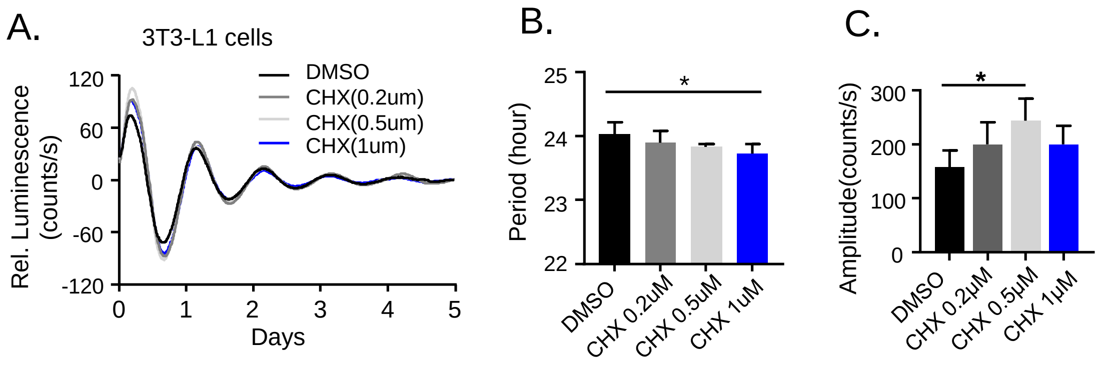


**Figure S1. Effect of chlorhexidine as a clock activator in 3T3-L1 preadipocytes.** (A) Baseline-adjusted tracing plots of average bioluminescence activity of Per2::dLuc reporter-containing 3T3-L1 preadipocytes (D-F) for 5 days, with quantitative analysis of clock period length (B) and cycling amplitude (C). Chlorhexidine (CHX) treatment was added to culture media at indicated concentrations. Data are presented as Mean ± SD of n=4 replicates for each concentration tested, for three independent repeat experiments.

#### Supplemental Figure S2.

A.

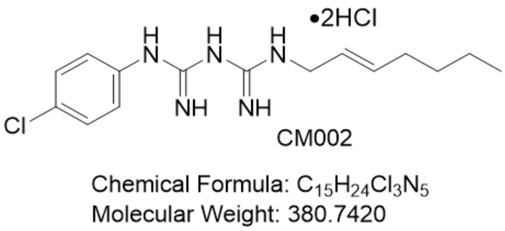


B.

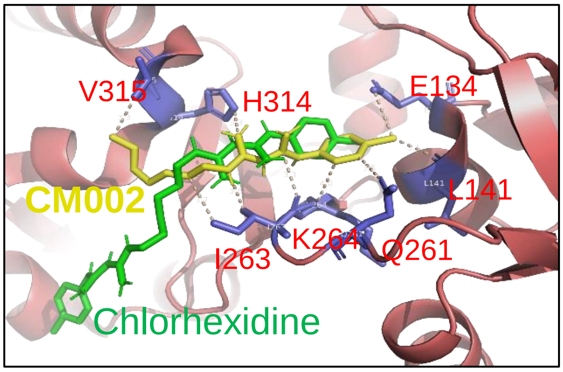


**Figure S2.** (A) Chemical structure of CM002. (B) Molecular docking modeling of CM002 (yellow) structure together with chlorhexidine (green) within the shared hydrophobic pocket of CLOCK protein (red). Predicted CM002 interactions with CLOCK protein residues were indicated.

#### Supplemental Figure S3.

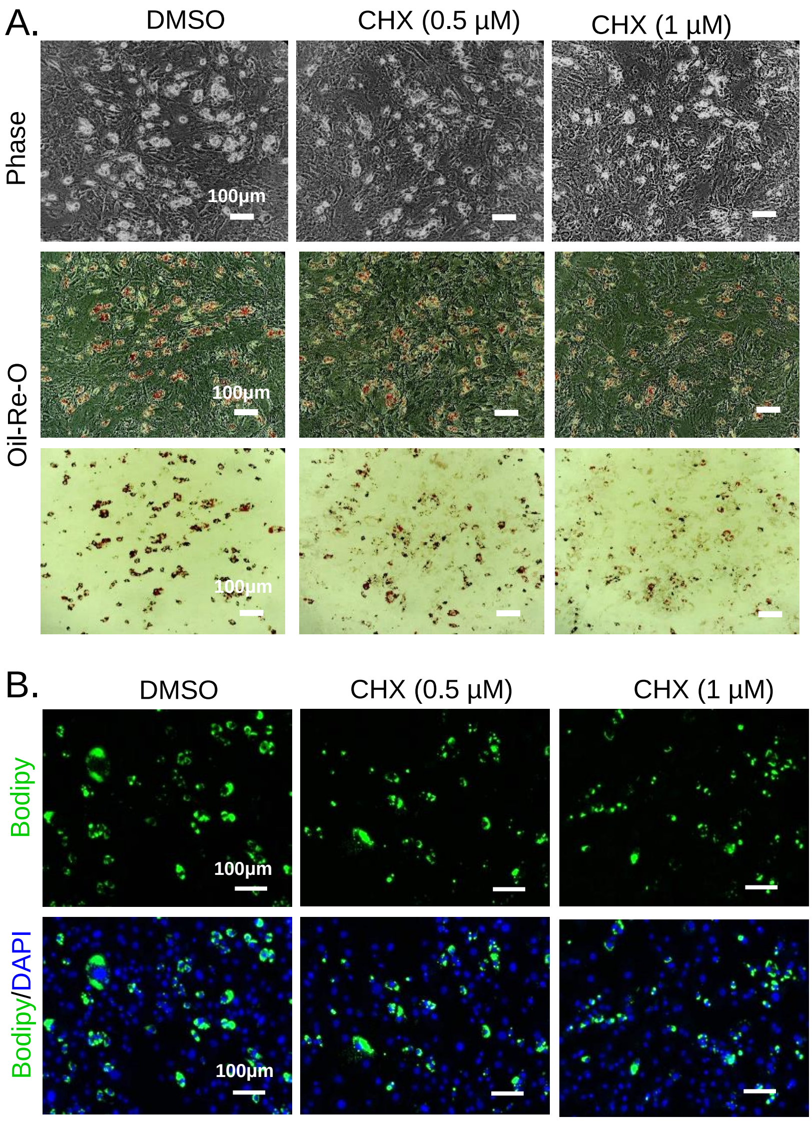


**Figure S3. Effect of Chlorhexidine on inhibiting early adipogenesis of primary preadipocytes.** (A) Representative images of phase-contrast and oil-red-O staining, and (B) Bodipy fluorescence staining of Primary preadipocyte at day 4 of early differentiation at indicated chlorhexidine (CHX) concentration. Scale bar: 100 µm.

#### Supplemental Table 1. Primary antibodies list.

| Antibody | Source | Cat# | Dilution |
|---|---|---|---|
| C/EBPα | Santa Cruz | SC-61 | 1:1000 |
| PPARγ | Santa Cruz | SC-7273 | 1:1000 |
| FASN | Santa Cruz | SC-48357 | 1:1000 |
| FABP4 | Santa Cruz | SC-271529 | 1:1000 |
| β-catenin | Cell Signaling | 8480S | 1:1000 |
| HSP90 | Cell Signaling | 4874S | 1:3000 |

#### Supplemental Table 2. Primer sequence for qPCR analysis.

| Genes | | Sequences |
|---|---|---|
| Bmal1 | Forward | CGCTTTCTGGAGGGTGTCCGC |
| | Reverse | TGCCAGGACGCGCTTGTACC |
| Clock | Forward | TTGCTCCACGGGAATCCTT |
| | Reverse | GGAGGGAAAGTGCTCTGTTGTAG |
| Nr1d1 | Forward | TGGCATCCGGTGCACTGCAG |
| | Reverse | CCCTCCAGAAGGGTAGCACGCT |
| Nr1d2 | Forward | GGAGTTCATGCTTGTGAAGGCTGT |
| | Reverse | CAGACACTTCTTAAAGCGGCACTG |
| Cry1 | Forward | CTGGCGTGGAAGTCATCGT |
| | Reverse | CTGTCCGCCATTGAGTTCTATG |
| Cry2 | Forward | TGTCCCTTCCTGTGTGGAAGA |
| | Reverse | GCTCCCAGCTTGGCTTGA |
| Per1 | Forward | CTGCCATGGAGGAAGAAGAG |
| | Reverse | AGCTGGGGCAGTTTCCTATT |
| Per2 | Forward | ATGCTCGCCATCCACAAGA |
| | Reverse | GCGGAATCGAATGGGAGAAT |
| Wnt1 | Forward | GGTTTCTACTACGTTGCTACTGG |
| | Reverse | GGAATCCGTCAACAGGTTCGT |
| Wnt10a | Forward | CAACGCGTGCGCTCTGGGTA |
| | Reverse | TGGCTCAAGCCCTTTCCGCG |
| Fzd2 | Forward | CATGCCCAACCTTCTTGGC |
| | Reverse | CAGCGGGTAGAACTGATGCAC |
| Fzd5 | Forward | AATCATGCAGGGGGCCCCGAA |
| | Reverse | CGACAAGCTAGGTACCTGTGGCG |
| Dvl2 | Forward | TCAGTTTGCGGGTGTGCGCAG |
| | Reverse | TTCGTCTCGCCTACACCACCG |
| Tcf4 | Forward | GGCCGCAGCGCCTTCTCTTTA |
| | Reverse | ACCATCATTGACTCCCCCGAGG |
| β-catenin | Forward | CGCTTGGCTGAACCATCAC |
| | Reverse | GTTCCGCGTCATCCTGATAGT |
| 36B4 | Forward | CGCTTTCTGGAGGGTGTCCGC |
| | Reverse | TGCCAGGACGCGCTTGTACC |
